## Supplementary material for "Using rare genetic mutations to revisit structural brain asymmetry": Supp. Fig.

### Supplementary Figures

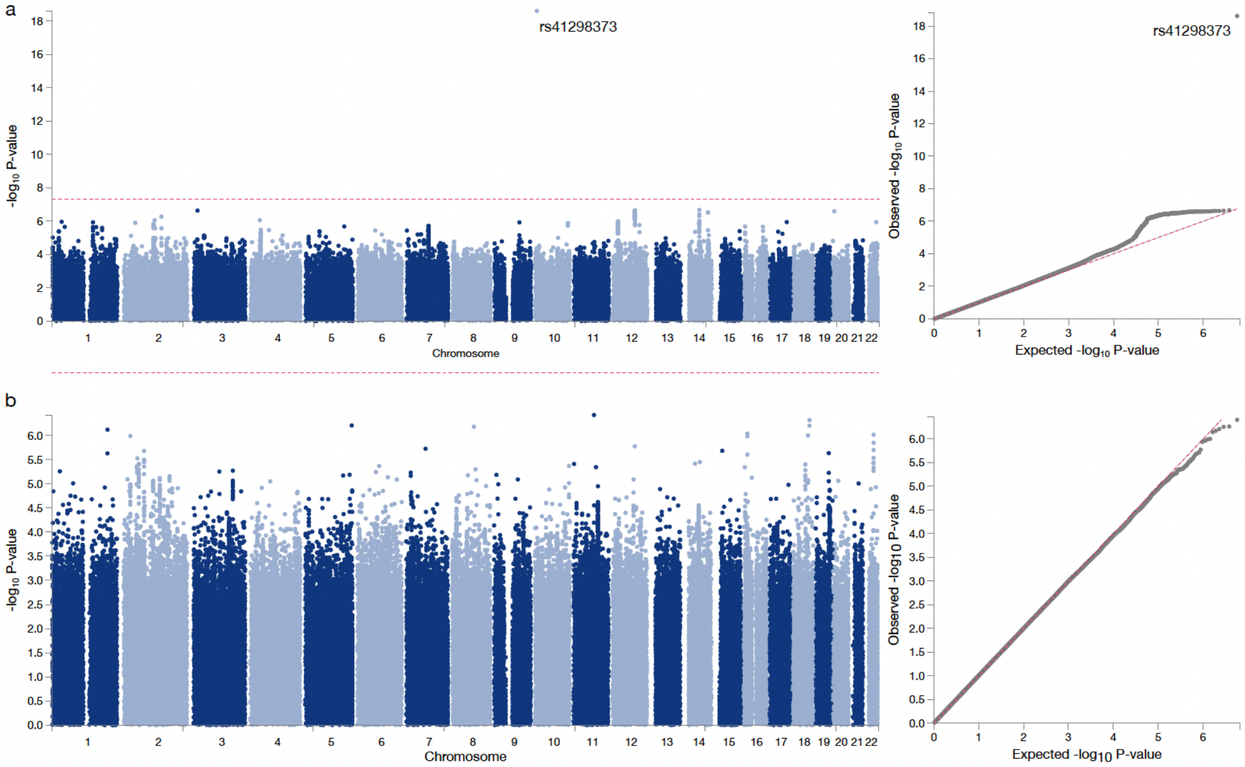


Supplementary Figure 1

**The influence of directionality of planum temporale asymmetry on GWAS results**

Our GWAS of planum temporale asymmetry revealed a single significantly associated SNP: rs41298373. Since planum temporale asymmetry is a directed measure, we investigated whether this directionality influences GWAS results. Specifically, we conducted two additional GWAS in the same pool of 30,358 UK Biobank subjects directed at a) the absolute value of planum temporale asymmetry and b) the absolute value of across-subjects z-scored planum temporale asymmetry. While computing an absolute value did not reveal additional SNPs, the use of an absolute z-score asymmetry index led to zero significantly associated SNPs. Therefore, different definitions of the asymmetry index did not highlight new genomic loci.

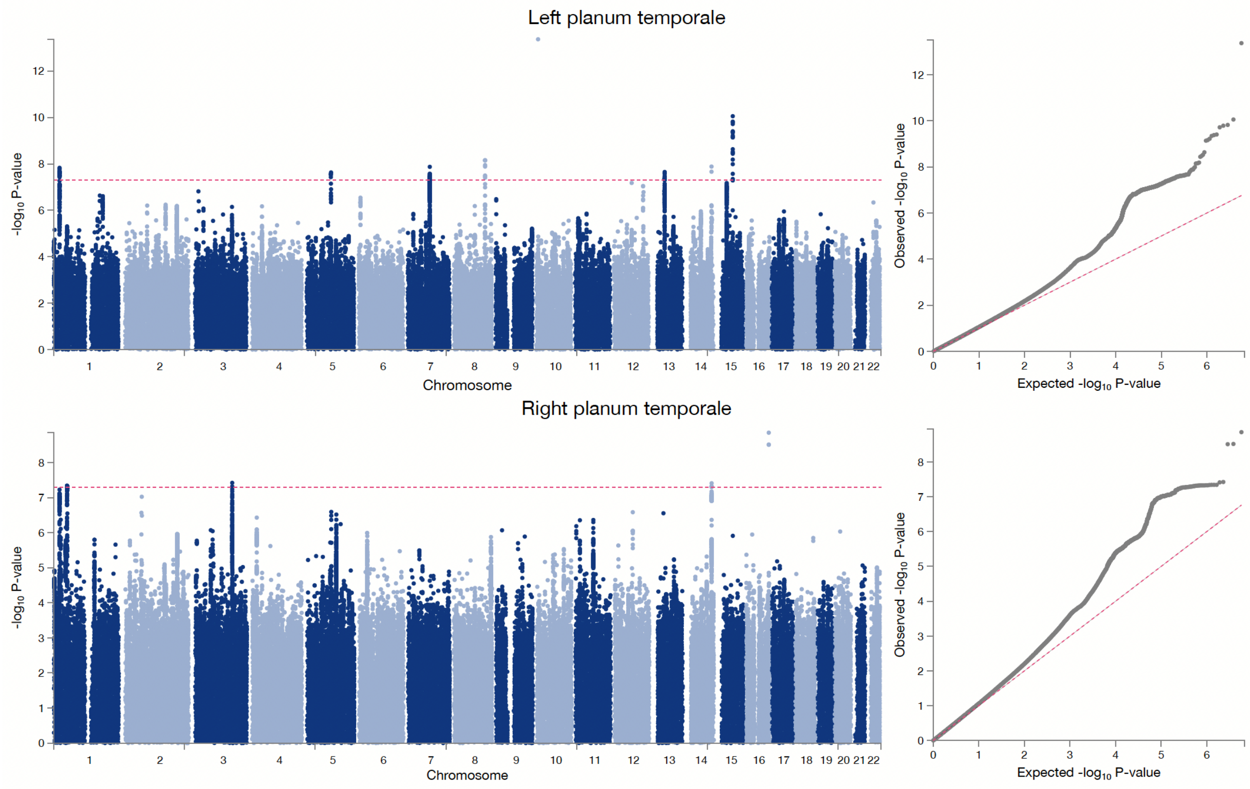


Supplementary Figure 2

**Dissecting the impact of common genetic variants on left and right planum temporale**

We conducted a GWAS (genome-wide association study) separately for the left and right planum temporale volume asymmetry in the 30,358 UK Biobank subjects. Based on GWAS of the left planum temporale volume, we found 726 significant candidate SNPs that mapped to eight genomic loci. Our GWAS of the right planum temporale volume yielded 368 candidate SNPs in four genomic loci. The QQ plots associated with both analyses suggest that genetic studies with more participants will likely locate additional loci.

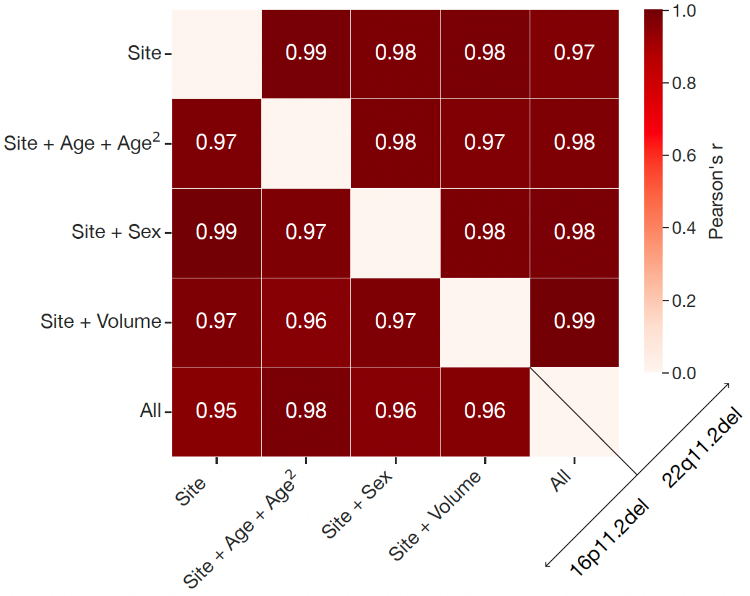


Supplementary Figure 3

**The effects of deconfounding factors on extracted patterns of brain asymmetry**

In our study, all derived regional brain volumes were adjusted for variation that can be explained by the scanning site. To probe the effect of other confounding variables, we further adjusted regional brain volumes for the effects of intracranial volume, age and age$^2$, sex, and all of these variables together. In the next step, we compared LDA asymmetry patterns derived using asymmetry indices based on these different preprocessing scenarios. We probed the influence of preprocessing on LDA models associated with both 16p11.2 and 22q11.2 deletion. Since all Pearson's correlations were > 0.95, adding more deconfounding variables did not influence the resulting asymmetry patterns for any CNV.

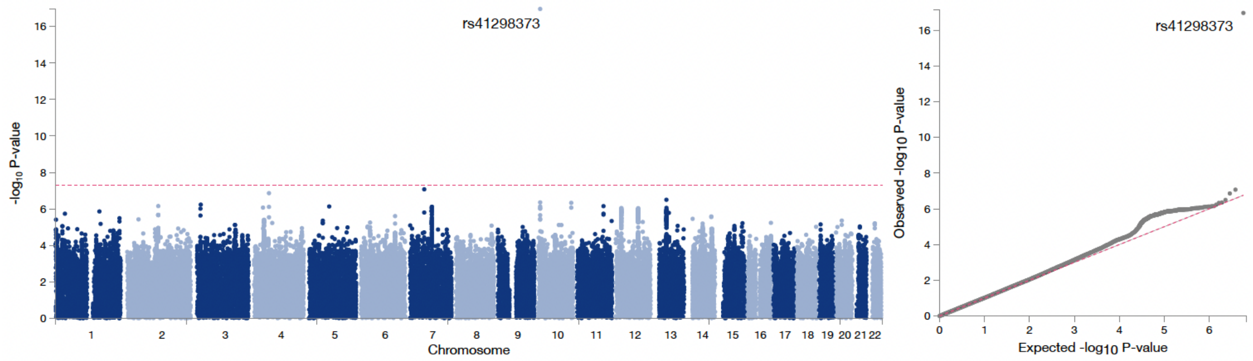


Supplementary Figure 4

**Genetic analysis of planum temporale with additional covariates**

To complement our original GWAS of planum temporale, we studied the influence of including total brain volume as an additional covariate. This preprocessing step did not lead to any change as the rs41298373 remained the only significantly associated SNP.
